# Coenzyme A homeostasis regulates hypoxic signalling via UBFD1 in triple-negative breast cancer

**DOI:** 10.1101/2025.06.18.660436

**Authors:** Paul Grevitt, Oscar Maiques, Kunal M. Shah, Valle Morales, Emanuela Gadaleta, Martin Dodel, Katiuscia Bianchi, Ivan Gout, J. Louise Jones, Claude Chelala, Michael J. Plevin, Faraz K. Mardakheh, Tyson V. Sharp

## Abstract

Hypoxia-inducible factors (HIFs) orchestrate cellular responses to oxygen deprivation and are frequently activated in triple-negative breast cancer (TNBC), driving metastasis and poor prognosis^1,2^. However, the full mechanisms of how HIFs are activated are not completely understood. It has been long noted that certain tumours have decreased levels of coenzyme A compared to surrounding tissues^3,4^, but the functional consequences of this remain unknown. Here, we identify Coenzyme A Synthase (CoAsy) as a novel regulator of HIF signalling. We demonstrate that CoAsy loss stabilises HIF-1α and HIF-2α independently of the canonical oxygen sensing pathway. Proximity-labelling proteomics revealed that CoAsy deficiency disrupts the association between HIF-1α and the proteasome through UBFD1, a novel CoA-binding protein that scaffolds HIFα for degradation. UBFD1’s CoA-dependent interaction with HIF-α is mediated by its PH domain, which undergoes CoAlation, an understudied post-translational modification. *COASY* loss of heterozygosity occurs in ∼1/3 of breast cancer patients and correlates with increased HIF activity, lack of hormone receptor expression and poor patient outcome. In vivo studies confirm that restoring CoAsy expression in tumours suppresses metastasis to the lung. These findings uncover a critical metabolic checkpoint regulating hypoxic signalling and identify CoAsy as a potential biomarker and therapeutic target in aggressive breast cancers.

## Main

The pseudohypoxic activation of hypoxia-inducible factors (HIFs) is a hallmark of many solid tumours, driving a cascade of metabolic and angiogenic adaptations that enhance tumour cell survival, proliferation and metastasis^5^. These processes are often linked to the dysregulation of key oncogenic pathways, including MYC^6^ and RAS^7^ hyperactivation and, most notably, the inactivation of the von Hippel-Lindau (VHL) tumour suppressor^8,9^. Furthermore, mutations in metabolic genes, such as fumarate hydratase (FH)^10^ in clear cell renal cell carcinoma (ccRCC), contribute to HIF stabilisation by increasing intracellular fumarate levels, which inhibit HIF-prolyl hydroxylases (PHDs), the enzymes responsible for initiating HIF degradation under normoxia ^11,12^. These metabolic and genetic aberrations illustrate the complex interplay between oxygen sensing and tumourigenesis.

Central to this regulation is the oxygen-sensitive hydroxylation of HIF-α subunits by PHD enzymes, with PHD2 (*EGLN1*) serving as the primary regulator. Recent evidence highlights a previously underappreciated mechanism of PHD2 inactivation under low intracellular cysteine levels, independent of oxygen availability^2^. This establishes a cysteine-sensing axis that amplifies HIF activity under specific metabolic conditions. Intriguingly, this mechanism is exploited in triple-negative breast cancer (TNBC), where tumour cells secrete glutamate to inhibit the cystine/glutamate antiporter (xCT), resulting in intracellular cysteine depletion and subsequent HIF stabilisation.

Despite the well-established pro-tumorigenic roles of HIFs in metabolic reprogramming, angiogenesis, and immune evasion, therapeutic targeting of HIFs in cancer has largely been confined to ccRCC, where HIF dependency is most pronounced^13^. Unlike other oncogenic drivers, such as mutant KRAS or MYC, HIFs exhibit a more context-dependent role in cancer progression, making their therapeutic exploitation challenging. Nevertheless, unravelling the nuanced regulatory networks governing HIF stability and activity may uncover novel, actionable vulnerabilities in cancers with limited treatment options. Here, we investigate an alternative regulatory axis involving coenzyme A (CoA) biosynthesis and its impact on HIF signalling, shedding light on a previously unrecognised metabolic checkpoint with broad implications for tumour biology.

### Coenzyme A Synthase (CoAsy) is a novel negative regulator of HIF signalling

To identify oxygen-independent regulators of HIF signalling, we transduced A549 cells with a hypoxia-responsive element (HRE) reporter (**Figure 1A)**, containing *firefly* luciferase under control of 8 canonical HREs (ACGTG). A549-HRE-*fluc* cells displayed increased luciferase activity upon exposure to hypoxia (1% O_2_) in a HIF-1α and time-dependent manner (**Figure S1A-D)**. We screened these lines against an arrayed siRNA library against all known kinases and pseudokinases in both normoxia and hypoxia (720 genes, **Figure S1E-F**). Hits with Z-scores ± 2 were used in a secondary validation screen, where we identified previous regulators of HIF activity including LATS2^14^ (**Figure 1B**). One consistent hit between normoxic and hypoxic screens was Coenzyme A synthase (CoAsy). CoAsy is a bifunctional enzyme that catalyses the final two steps of *de novo* CoA biosynthesis from pantothenic acid (Vitamin B5)^15^. As CoA is membrane impermeable, this is the sole pathway for cells to synthesise CoA. Extracellular 4’-phosphopantetheine can also act as a substrate for CoA biosynthesis, yet still requires CoAsy activity to produce mature CoA^16^. Due to the importance of this pathway for life, we questioned whether cells would be viable with decreased levels of CoAsy; to address this, we consulted the DEPMAP dataset^17^. *COASY* scored as an essential gene with negative CERES scores (CRISPR screens), but did not rank as an essential gene in RNAi screens (DEMETER score); this contrasts with the essential gene *RPL5*, which ranked as essential in both RNAi and CRISPR screens (**Figure S1G**). This suggests that cells can tolerate low levels of CoAsy, but not complete loss, and thus may indeed be a mechanism contributing to HIF biology.

**Figure 1:**
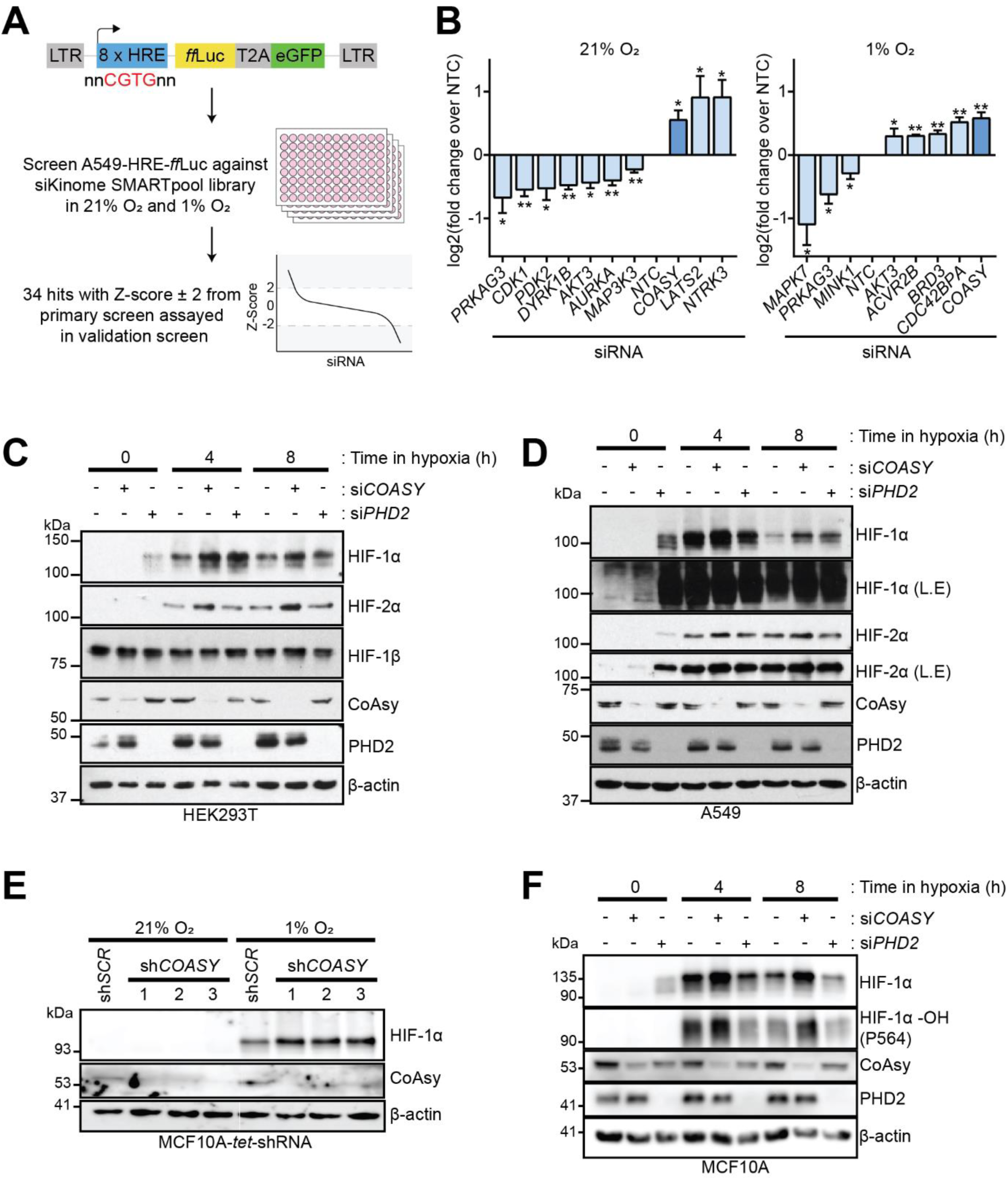
RNAi screen identifies Coenzyme A Synthase as a novel negative regulator of HIF1 signalling. **A)** Schematic of HIF-reporter screen. **B)** Significant hits from validation screen in normoxia and hypoxia (n = 3). **C)** siRNA knockdown of *COASY* in HEK293T cells cultured in normoxia or hypoxia (n = 3). **D)** siRNA knockdown of *COASY* in A549 cells cultured in normoxia or hypoxia (n = 3). **E**) Doxycycline inducible shRNA knockdown of *COASY* in MCF10A cells cultured in normoxia or hypoxia (4 h) (n = 3). **F)** siRNA knockdown of *COASY* in MCF10A cells cultured in normoxia or hypoxia (n = 3). L.E = long exposure. * p < 0.05, ** p < 0.01, *** p < 0.001

RNAi knockdown of *COASY* revealed an increase in both HIF-1α and HIF-2α protein levels over scrambled control, but no difference in HIF-1β suggesting that this is a HIFα-specific effect (**Figure 1C-D**). This was validated using distinct RNA sequences from the primary screen across a range of cell lines from differing tissues of origin (osteosarcoma, lung, breast) (**Figure 1C-F, Figure S1J-K**). This includes non-tumorigenic MCF10A cells (**Figure 1E-F**), highlighting that loss of CoAsy alone is sufficient for increases in HIFα, and does not require other oncogenic drivers. Importantly, there was no decrease in levels of hydroxylated HIF-1α (P564), indicating that the primary oxygen sensing pathway (hydroxylation by HIF-prolyl hydroxylases) is not inhibited upon loss of CoAsy (**Figure 1F**).

### CoA biosynthesis influences HIFα protein levels via the proteasome independently of PHD-VHL axis

The rate limiting step in CoA biosynthesis occurs at the initial step, through phosphorylation of pantothenic acid by pantothenate kinases (PANKs)^18^, these are regulated by acyl-CoAs (**Figure 2A**). However, the beta isoform of PANK1 is less susceptible to this negative feedback loop, therefore overexpression of this isoform increases levels of CoA and CoA derivatives^19^. Cells expressing this isoform have significantly reduced levels of both HIF-1α and HIF-2α compared to control (**Figure 2B**). Furthermore, MCF10A (**Figure 2C**) and triple-negative breast cancer cell lines (**Figure S2A-B**) overexpressing Pank1β or CoAsy display reduced HIF-1α compared to controls. This suggests that CoAsy regulates HIF via catalysing CoA biosynthesis as opposed to a non-canonical function of the CoAsy protein. To further evaluate this, we cultured Hs578t-HRE-*fluc* cells in media with decreasing concentrations of calcium pantothenate, starting from 4 mg.L-^1^, the concentration found in basal DMEM medium. Notably, cells cultured in calcium pantothenate-depleted medium exhibited increased HRE-reporter activity compared to those maintained at 4 mg.L^-1^ (**Figure 2D**).

**Figure 2:**
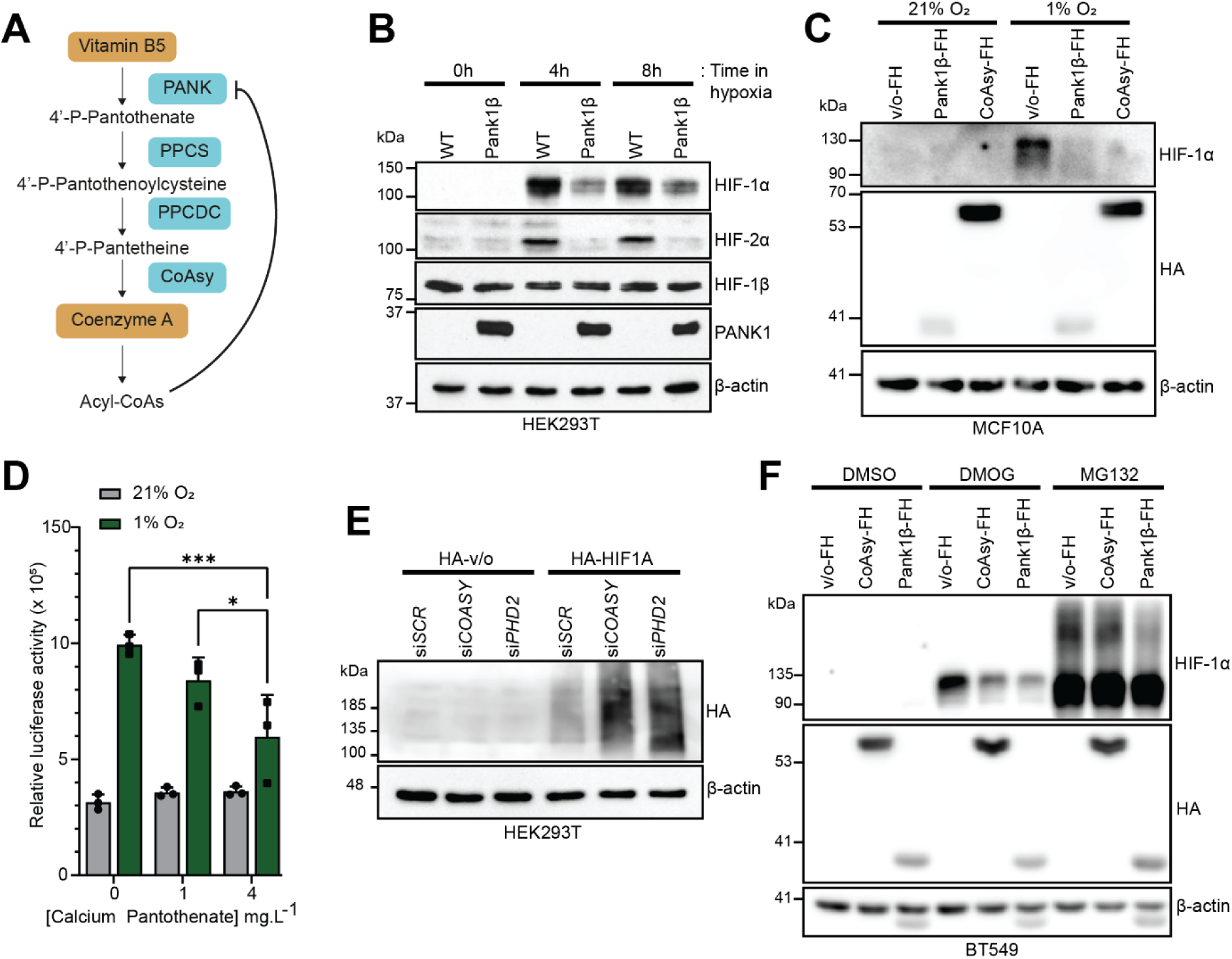
Coenzyme A biosynthesis is coupled with HIF signalling. **A**) Schematic of de novo CoA biosynthesis from Pantothenate (Vitamin B5). **B)** HEK293T cells overexpressing Pank1β in normoxia and hypoxia (n = 3). **C)** Overexpression of Pank1β-FlagHA or CoAsy-FlagHA in normoxia or hypoxia (4 h) (n = 3). **D)** HRE-reporter activity in Hs578t cells cultured in varying concentrations of calcium pantothenate in normoxia and hypoxia (representative graph from n = 2). **E)** Overexpression of HA-HIF1A in HEK293T cells with *COASY* knocked down via siRNA (n = 3). **F)** Treatment of B549 CoAsy-FlagHA or Pank1β-FlagHA cells with DMOG (50 μM for 16h) or MG132 (10 μM for 2h) (n = 3). * p < 0.05,** p < 0.01, *** p < 0.001.

We did not observe any differences in *HIF1A* or *EPAS* (encodes HIF-2α) following knockdown of *COASY* (**Figure S2C**), therefore, we sought to examine if loss of CoAsy effects HIFα post-transcriptionally. We expressed an ectopic HA-HIF1A construct, with synthetic 5’ and 3’ UTRs to negate endogenous transcriptional and translational regulatory mechanisms, in cells with si*COASY.* Knockdown of *COASY* resulted in increased HA-HIF1A compared to scrambled control, in line with positive control si*PHD2* (**Figure 2E**). We treated CoAsy or Pank1β overexpressing cells with a pan-dioxygenase inhibitor (DMOG) or MG132 to inhibit the proteasome. DMOG treatment blocks prolyl hydroxylation of HIFα by the PHDs, which in turn stops pVHL binding, shutting down the canonical oxygen-dependent pathway of HIF regulation. DMOG treatment stabilised HIF-1α under control conditions; however, this stabilisation was reduced in cells overexpressing CoAsy or Pank1β (**Figure 2F**). In contrast, treatment with MG132 stabilised HIF-1α to the same extent across all cell lines, indicating that CoA biosynthesis regulates HIFα via the proteasome independently of the PHD-VHL axis.

### UBFD1 is a CoAsy-dependent interactor of HIF-1α

Given that our data indicated CoA biosynthesis modulates HIFα stability through a non-canonical proteasome pathway, we aimed to identify novel interactors of HIF-1 that are altered upon the loss of *COASY.* To achieve this, we utilised a proximity labelling approach, fusing a biotin ligase (BioID2^20^) to HIF-1α and expressing this in sh*SCR* or sh*COASY* cells treated with MG132. Through this approach, we identified proteins that were proximal to HIF-1α, with 497 identified in sh*SCR* and 675 in sh*COASY* cells (**Figure 3A-B**). This analysis revealed several known interactors, including ARNT (HIF-1β) and STAT3^21^, both of which showed increased association following the loss of *COASY*. In addition, we observed a decrease in association with components of the proteasome in sh*COASY* cells, which supports the notion that CoAsy regulates HIF via the ubiquitin proteasome system (UPS) (**Figure 3C**). To identify the mechanism by which this is occurring, we identified proteins within the UPS and compared between sh*SCR* and sh*COASY* conditions (**Figure 3D**). This revealed Ubiquitin Family domain-containing 1 (UBFD1) as the sole difference between conditions, with association detected in sh*SCR* conditions but not in sh*COASY* (**Figure 3E-F**). Little is known about the function of UBFD1, other than a reported ability of binding polyubiquitin chains^22^; the structure prediction by alphafold2 identifies the ubiquitin-like domain as well as an unstructured N-terminus and a C-terminal PH domain (**Figure 3G**). We confirmed that HIF-1α and UBFD1 interact in an overexpressed IP (**Figure S3A**), with further evidence of altered association between HA-HIF1A and endogenous UBFD1 in sh*SCR* and sh*COASY* cells (**Figure 3H**). These findings highlight a CoA-dependent regulatory mechanism involving UBFD1, which may modulate HIF stability through specific protein-protein interactions.

**Figure 3:**
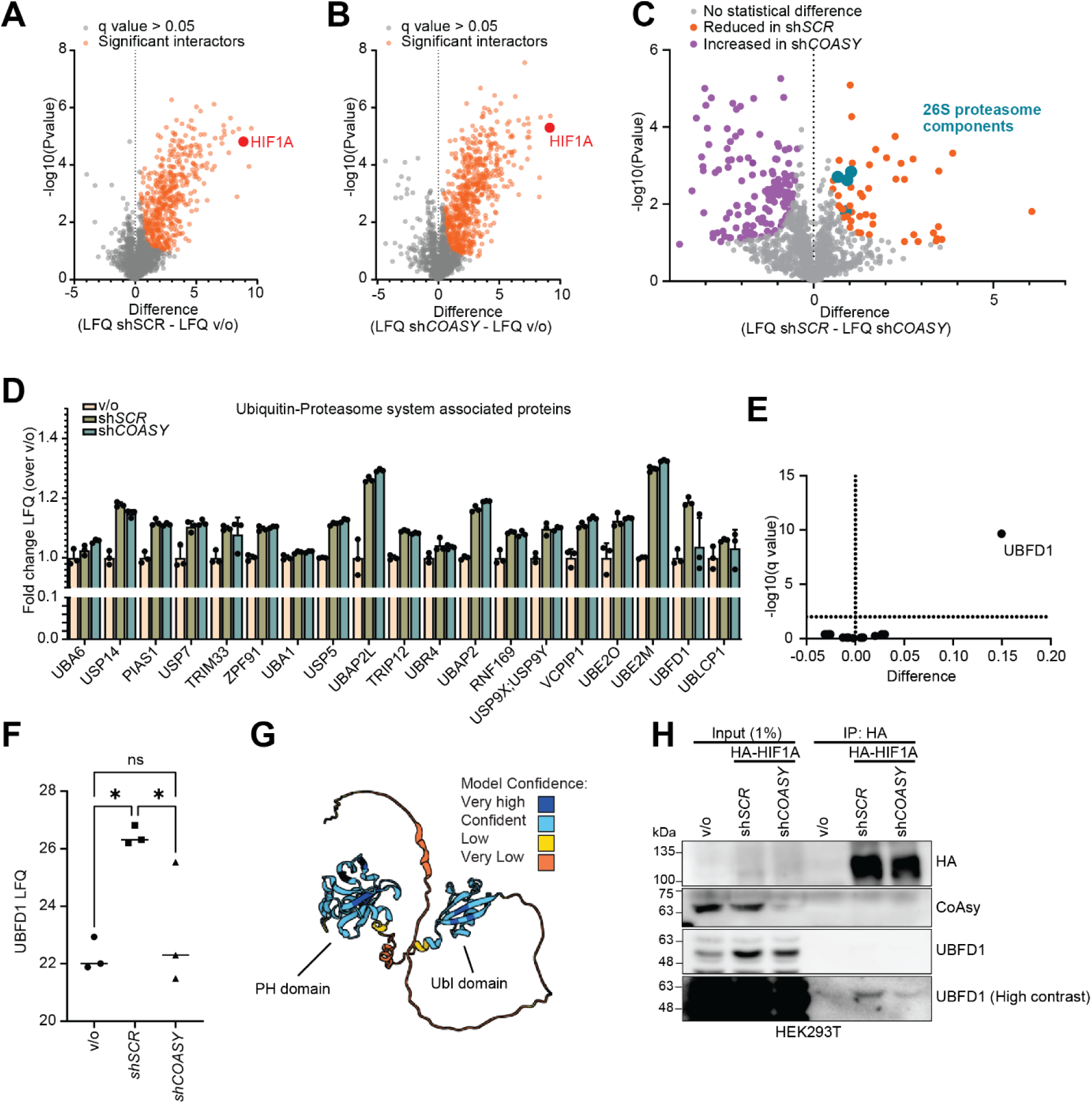
Proximity labelling-MS identifies UBFD1 as a CoAsy dependent interactor of HIF-1α. **A-B)** Volcano plot of proteins enriched in HIF-1A BioID2 MS in HEK293T sh*SCR* (**A**) and sh*COASY* (**B**) compared to vector only cells. **C)** Volcano plot of proteins differentially enriched in sh*COASY* compared to sh*SCR* cells. **D**) Bar chart of ubiquitin-proteasome system (UPS) associated proteins identified in HIF-1A BioID2 MS. **E**) Volcano plot of UPS proteins differentially expressed in sh*SCR* and sh*COASY* cells. **F)** Label-free quantification of UBFD1 from HIF-1A BioID2 MS. **G)** Predicted structure of UBFD1 from alphafold server. **H**) Pulldown of HA-HIF1A in sh*SCR* and sh*COASY* cells (n = 1). * p < 0.05, ** p < 0.01, *** p < 0.001.

### UBFD1 is a CoA sensor and controls association of HIFα with the proteasome

Mapping of the interaction between HIF-1α and UBFD1, reveals that UBFD1 binds via the cryptic PH domain (**Figure 4A**). As the binding of UBFD1 to HIF-1α is dependent on the PH domain and CoA availability, we postulated whether UBFD1 could act as a CoA sensor. We observed binding of UBFD1 to CoA agarose beads, which is lost upon deletion of the PH domain (**Figure 4B**). Building on the evidence that UBFD1 interacts with HIFα in a CoA-dependent manner, we sought to investigate whether CoA or acetyl-CoA could enhance this interaction. To test this, we incubated an IP reaction of HA-HIF1A and Flag-UBFD1 with CoA or acetyl-CoA. Here, we observed increased association when incubated with CoA and to a lesser extent with acetyl-CoA, suggesting that this interaction is partially dependent on CoA. (**Figure 4C**). Furthermore, the melting temperature (*T*_m_) of the PH domain increased upon addition of CoA, which is consistent with stabilisation by ligand binding. By contrast, no change with pantothenate (**Figure 4D-E**). This change in protein stability was lost at low concentrations of CoA (40 μM) which is below the reported cytoplasmic concentration for CoA^23,24^ (**Figure S4A**). Recombinant PH domain bound to CoA agarose beads; however, it could not be eluted in the presence of supraphysiological levels of CoA, whereas was eluted under high SDS and reducing conditions (eluted from beads in Laemmli buffer) suggesting covalent modification (**Figure S4B**). CoAlation is a recently described post-translational modification whereby a thiol linker is formed between free cysteine residues and CoA^19^. We confirmed CoAlation of recombinant PH domain with an anti-CoA antibody (**Figure 4F**), and that this occurred at the sole cysteine within this domain (**Figure S4C**). Full length UBFD1 could be CoAlated in cellulo, however we did not see a complete ablation of CoAlation upon mutating C185A, suggesting that the cysteine in the unstructured N-terminus of UBFD1 can also be CoAlated (**Figure S4D**).

**Figure 4:**
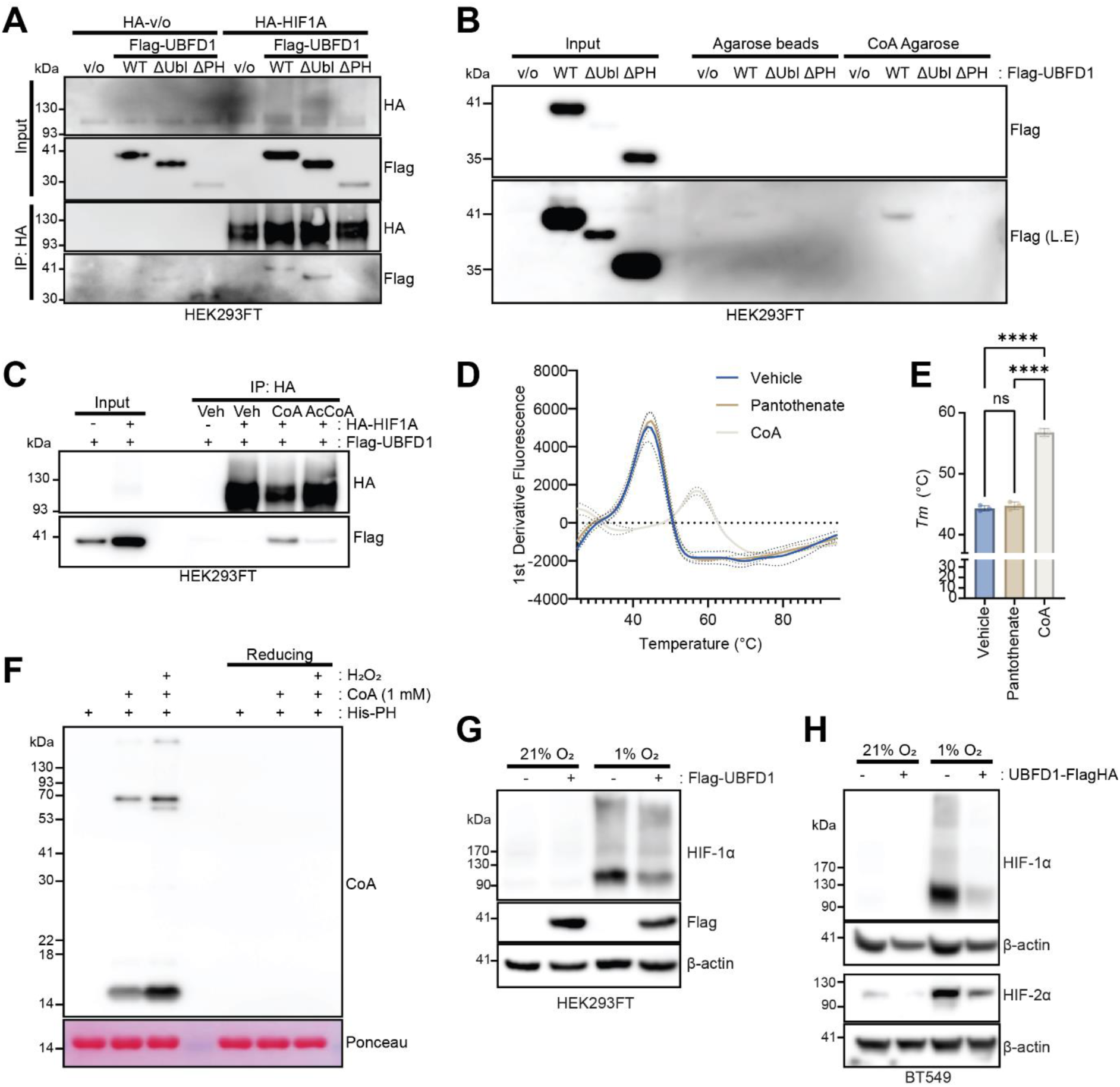
UBFD1 is CoAlated which controls association with HIF-1α. **A)** Overexpressed co-immunoprecipitation of HA-HIF1A in HEK293FT cells with Flag-UBFD1 or domain deletion mutants (n = 3). **B)** Pulldown of Flag-UBFD1 or domain deletion mutants with CoA-Agarose beads (n = 3). **C)** Co-immunoprecipitation of HA-HIF1A with Flag-UBFD1 in the presence of CoA or Acetyl-CoA (150 μM) (n = 3). **D-E)** Thermal shift assay of His-UBFD1 PH domain with pantothenate or CoA (1 mM) (n = 3). **F)** Western blot from in vitro CoAlation of His-UBFD1-PH domain. **G)** Overexpression of Flag-UBFD1 in HEK293FT cells cultured in normoxia or hypoxia (4 h) (n = 3). **H)** Lentiviral overexpression of UBFD1-FlagHA in BT549 cells cultured in normoxia and hypoxia (4 h) (n = 3). * < 0.05, ** p < 0.01, *** p < 0.001, **** p < 0.0001.

Overexpression of UBFD1 reduces both HIF-1α and HIF-2α in normoxia and hypoxia (**Figure 4G-H**). Next, we sought to determine how UBFD1 was regulating HIF protein levels. We observed an increased association between 19S proteasome components (PSMD2 and PSMC1) with HA-HIF1A upon overexpression of Flag-UBFD1 (**Fig S4E**). In addition, we performed an endogenous IP of HIF-1α in MG132 treated MB231 cells with UBFD1 knocked out via CRISPR-Cas9. Here we observed a reduced association with both 19S and 20S proteasome components, indicating that UBFD1 is acting as a scaffold of HIF-1α to the proteasome (**Figure S4F**). Collectively, these findings demonstrate that UBFD1 functions as a CoA-binding and sensing protein, modulating HIFα degradation via the proteasome in a CoA-dependent manner.

### CoAsy LOH is a prognostic marker in breast cancer

To explore the clinical relevance of CoAsy loss, we examined the copy number alterations in 3 breast cancer datasets and observed shallow deletions (representing loss of one allele) of *COASY* in 27-35% of patients (**Figure 5A**). We did not observe a significant number of deep deletions, which supports our hypothesis that cells can tolerate low levels of COASY, but not complete loss. In our in-house cohort of breast cancer patients^25^, we compared *COASY* expression between primary tumour to histologically normal adjacent tissue (<2 cm from primary tumour) and surrounding tissue (>5 cm from primary tumour) and observed a significant reduction of *COASY* expression in tumour compared to non-cancerous tissue (**Figure 5B**). We further stratified patients in TCGA and METABRIC datasets and observed lower *COASY* expression in ER and PR negative patients (**Figure 5C-D**) which was also observed in CPTAC proteomics dataset (**Figure 5E**). This trend was confirmed in a panel of breast cancer lines, where the triple-negative breast cancer lines had lower levels of CoAsy, compared to receptor positive (**Figure 5F**). We correlated *COASY* mRNA expression with genes in the Buffa hypoxic signature^26^ and observed a negative correlation with the majority of these in METABRIC and TCGA datasets, indicating the relationship between CoAsy and HIF signalling we observe in vitro is also occurring in the tumours of breast cancer patients. Importantly, we observed that reduced *COASY* expression or copy number alterations correlated with poor patient outcome (**Figure 5G**). These data show that *COASY* is lost frequently in breast cancer and correlates with poor patient outcomes.

**Figure 5:**
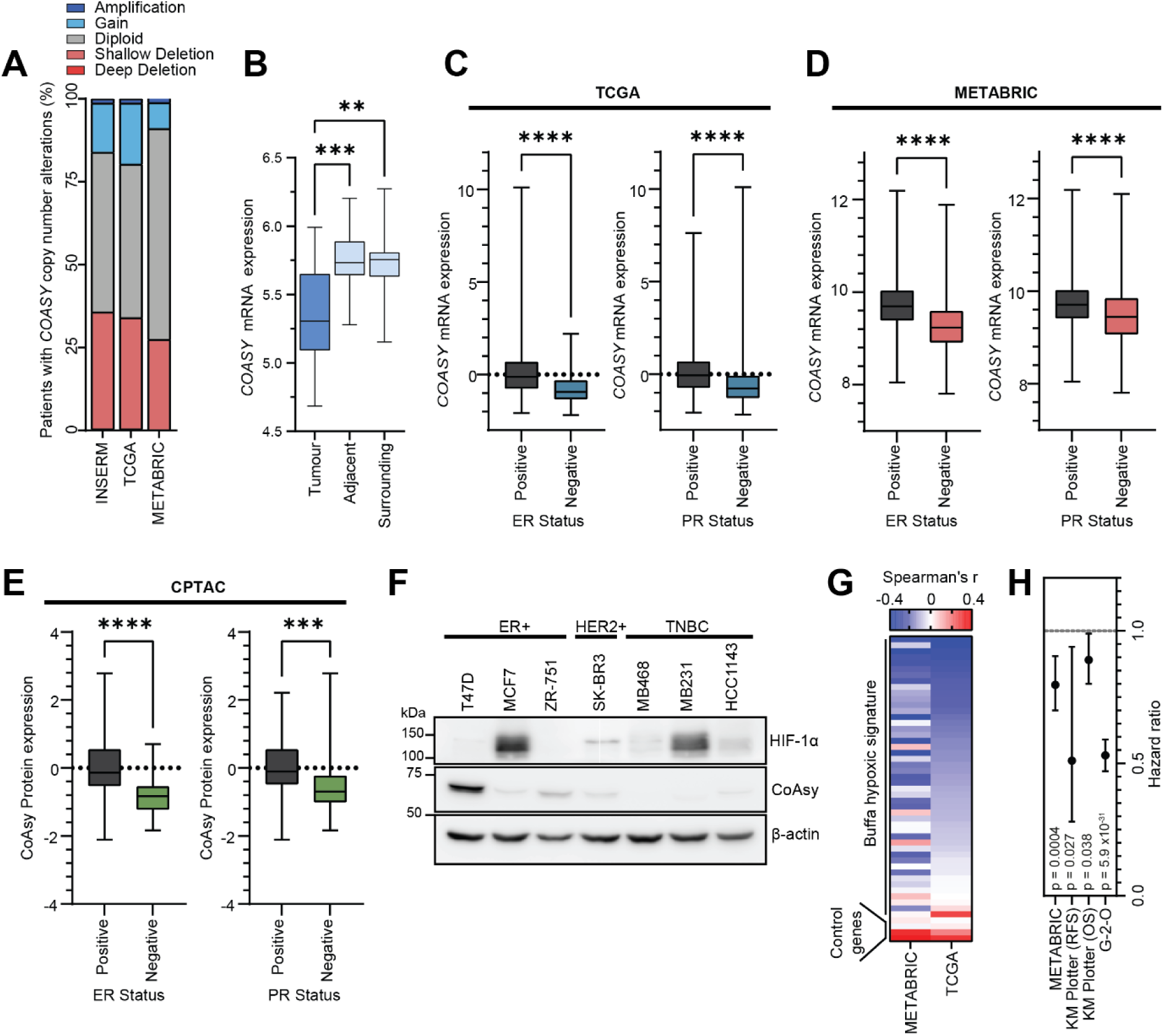
*COASY* LOH occurs in breast cancer and correlates with receptor status and prognosis. **A)** Putative *COASY* copy number alterations from 3 patient cohorts. **B)** *COASY* mRNA expression from in-house cohort of 19 patient’s tumour, adjacent tissue (<2 cm cm from tumour) or surrounding tissue (>5 cm from tumour). **C)** *COASY* mRNA expression in patients from TCGA cohort stratified by hormone receptor status. **D)** *COASY mRNA* expression in patients from METABRIC cohort stratified by hormone receptor status. **E)** CoAsy protein levels from CPTAC consortium stratified by hormone receptor status. **F)** Western blot of CoAsy protein expression in panel of breast cancer cell lines (n = 3). **G)** Heatmap of Spearman correlation coefficients of *COASY* mRNA expression and genes in Buffa hypoxic signature in TCGA and METABRIC cohorts. **H)** Plot of Hazard ratios of patient survival stratified by *COASY* expression. * < 0.05, ** p < 0.01, *** p < 0.001, **** p < 0.0001.

We examined other cancer types, such as lung cancer where we did not see the same degree of LOH and a positive correlation with genes in Buffa hypoxic signature, suggesting there may be some cancer type selectivity (**Figure S5A-B**). Interestingly, we observed a decrease of *COASY* and other genes (*PANK1, PANK3* and *PPCS*) from the CoA biosynthesis pathway in ccRCC tumours compared to normal tissue (**Figure S5C**). Intriguingly, low expression of pantothenate kinase isoforms correlates with poor patient outcome; conversely, high levels of the terminal step of this pathway, *COASY* displays poor patient outcome (**Figure S5D-F**). It is unclear as to the significance of this divergence between genes in this pathway, potentially suggesting other factors such as moonlighting functions of CoAsy, including its interaction with p85α^27^ or S6K1^28^, could have secondary effects. Notably, previously published comparative metabolomics data in ccRCC patients^29^, showed reduced CoA levels in primary tumours compared to adjacent normal tissue, as well as reduced levels of acetyl CoA and pantothenate, indicating that Vitamin B5 import may also be decreased in ccRCC (**Figure S5G**).

### CoAsy controls breast cancer metastasis

To assess the consequences of increased expression of genes in CoA biosynthesis pathway in TNBC cell proliferation, we performed colony formation assays in HCC1143 and BT549 cells overexpressing CoAsy or Pank1β in normoxia and hypoxia. In both normoxia and hypoxia we observed a decrease in colony forming ability indicating reduced cell fitness upon CoAsy or Pank1β overexpression (**Figure 6A-B**). We further tested cell fitness in MB231 cells overexpressing CoAsy or a CoAsy mutant (R499C), found in rare cases of neuronal brain iron accumulation (NBIA) disorder termed CoPan that results in loss of Dephospho-CoA kinase activity, where patients have reduced levels of mature CoA and acyl-derivatives^30^. Here, we observed a decrease in colony forming ability upon overexpression of WT CoAsy but no difference upon overexpression of CoAsy R499C mutant (**Figure 6C**). Next, we sought to examine the effects of restoring CoAsy function in TNBC lines in vivo. To achieve this, we implanted MB231 cells overexpressing WT CoAsy or R499C mutant in the mammary fat pad of Nod Scid gamma (NSG) mice. Perhaps surprisingly, we did not observe differences in primary tumour growth between the lines (**Figure 6D-E**), with minimal differences in tumour necrosis, apoptosis or proliferation (**Figure S6A-C**). However, we observed a significant decrease in lung metastasis upon overexpression of WT CoAsy compared to vector only control, as evidenced by decrease in total lung weight and reduced percentage of tumour staining in lungs indicating a reduced metastatic burden (**Figure 6F-H**).

**Figure 6:**
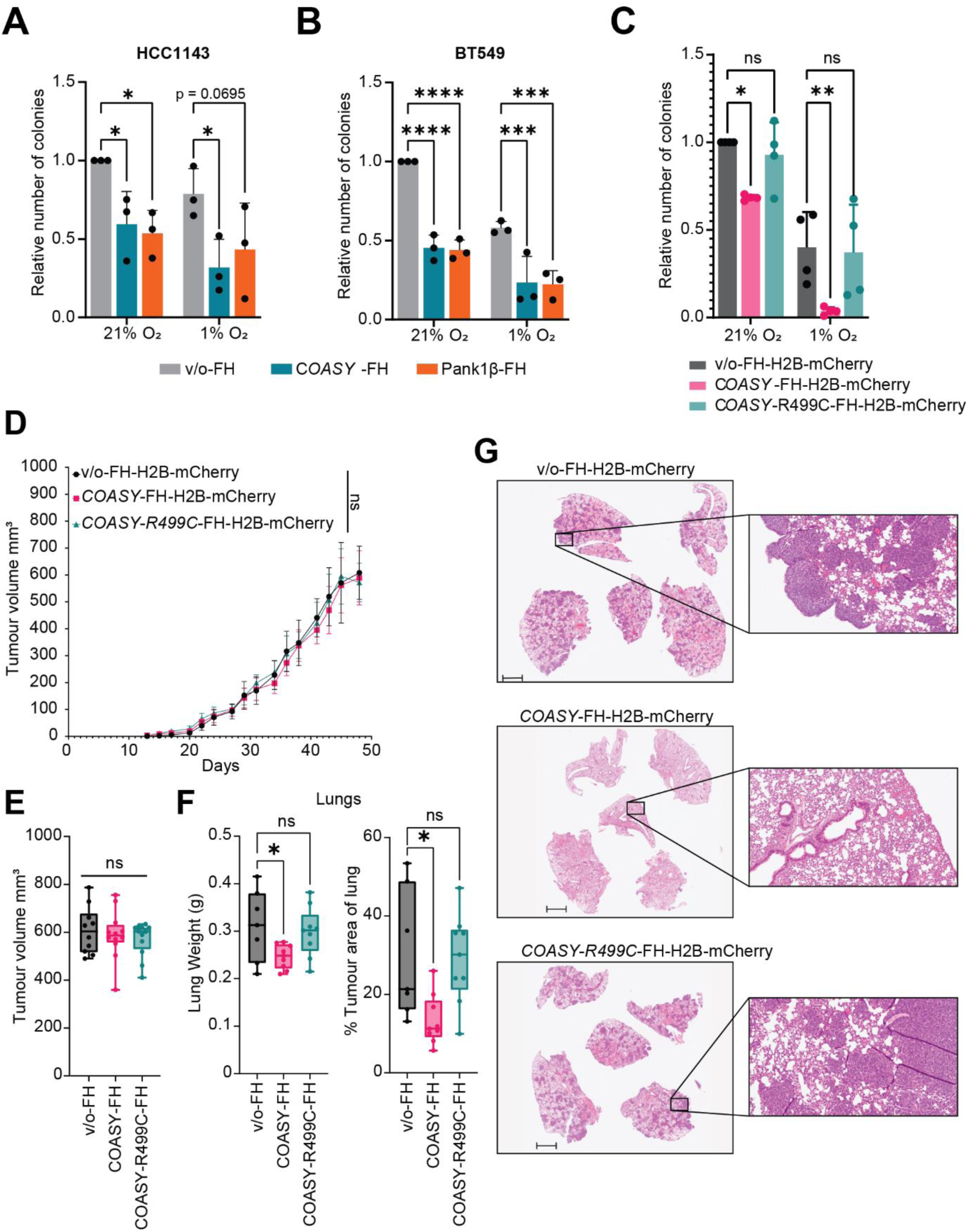
CoAsy overexpression reduces breast cancer metastasis. **A-B)** Colony formation assay of HCC1143 or BT549 cells overexpressing CoAsy-FlagHA or Pank1β-FlagHA in normoxia or hypoxia (n = 3). **C)** Colony formation assay of MB231 overexpressing CoAsy-FlagHA or CoAsy R499C mutant in normoxia or hypoxia (n = 4). **D)** Orthotopic tumour growth of MB231 cells overexpressing CoAsy-FlagHA or CoAsy R499C mutant in NSG mice (vector only n = 7, CoAsy-FlagHA and CoAsy R499C-FlagHA n = 9). **E)** Final tumour measurements from orthotopic xenograft study. **F)** Weight of lungs and percentage of lung area as tumour from mice in orthotopic xenograft study. **G)** Representative H&E of lungs from mice in orthotopic xenograft study. * < 0.05, ** p < 0.01, *** p < 0.001

## Discussion

Here we present Coenzyme A Synthase as a novel regulator of HIFα stability in breast cancer, and a coupling of CoA homeostasis to hypoxic signalling. The protein UBFD1 undergoes covalent modification by CoA which promotes interaction with HIF-1α, promoting association with the proteasome, increasing its degradation. These findings provide a mechanism by which cells can sense changes in levels of CoA, with reduced CoAlation of UBFD1 below cytoplasmic concentrations, resulting in stabilised HIF and increased signalling. Studies have identified tumours with reduced levels of CoA compared to adjacent normal tissue^3,4^, however until now it has been unclear as to why this was occurring. When one would posit that increased CoA would allow for heightened metabolic activity and increases in other CoA dependent reactions such as histone acetylation, both required for proliferating cancer cells. Our data provides a potential rationale for this, with TNBC cells co-opting this mechanism through *COASY* LOH to activate HIF and rewire the metabolism to maintain proliferation, growth and metastasis. Whilst these data indicate that UBFD1 can regulate HIF signalling via CoAlation at its cryptic PH domain, it remains unknown whether other proteins are susceptible to regulation via this mechanism, and the impact that these have on cellular transformation.

Our data highlights that this occurs frequently in breast cancer, and there is a degree of cancer type specificity between the regulation of HIF by CoAsy. Indeed, there may exist an additional layer of cell type specificity. IL-22 producing CD8+ T cells have increased CoA biosynthesis^31^, which in contrast to our findings in breast cancer cells, display increased HIF-1α. However, increased supplementation of CoA, increased T-cell killing of tumours; thereby suggesting that CoA supplementation may be of benefit to both cell intrinsic and extrinsic mechanisms in breast cancer. In myelodysplastic syndromes, characterised by mutant SF3B1, there are reduced levels of CoAsy because of altered splice variants^32^. Importantly, observed erythropoiesis defects can be reversed upon supplementation with Vitamin B5. Furthermore, recent studies have shown that supplementing mice with exogenous CoA reduced tumour volume in the PyMT mouse model of breast cancer and increased α-CTLA4 efficacy, in part due to increased macrophage anti-tumour activity^33^. Further supporting our hypothesis that supplementation of Vitamin B5 or CoA could enhance breast cancer treatment.

Despite improvements in immunotherapy, there is still a need for targeted therapies in TNBC. With *COASY* LOH observed in ∼1/3 of breast cancer patients, and significant overlap with receptor negative patients, a targeted therapy against CoAsy deficient cells may be of clinical benefit. It has been reported that reduction of CoA can increase sensitivity to ferroptosis^34^ and *COASY* has scored highly in screens for sensitivity to ferroptosis inducing agents (FINs)^35^, therefore *COASY* LOH may be a potential biomarker for the next generation of FINs. We observed a decrease in lung metastasis when CoAsy function was restored to TNBC cells. Activation of HIFs has been linked to promoting metastasis through promoting EMT via upregulation of target genes such as *TWIST1*^36^. With secondary effects arising due to metastatic burden being the primary cause of breast cancer mortality, targeting CoAsy deficiency is therefore an attractive therapeutic avenue. This study highlights a previously unrecognized CoA-dependent regulatory axis in cancer biology, offering not only a deeper understanding of metastasis but also a promising avenue for the development of targeted therapies in aggressive breast cancers.

## Methods

### Cell Culture

Cells were maintained in DMEM (A549, HEK293T, HEK293FT, U2OS, MDA-MB231, MDA-MB436, MDA-MB468 & Hs578t), RPMI (BT549, T47D, MCF7, ZR-751 HCC1143) or McCoy’s 5A (SK-BR3) supplemented with 10% FBS and 1% penicillin-streptomycin. MCF10A cells were cultured in HUMEC media (Gibco, 12752010). All cells were maintained in a humidified 37⁰C incubator with 5% CO_2_. For low pantothenate medium, basal DMEM without glucose, L-glutamine, D-pantothenic acid, pyruvic acid (US biological life sciences, D9800-02B) was supplemented with glucose, GlutaMAX, sodium pyruvate and sodium bicarbonate before supplementing with 10% charcoal stripped FBS (Sigma-Aldrich, F6765) and 1% penicillin-streptomycin. All cell cultures were regularly tested for mycoplasma. Parental stocks of each line were obtained from ATCC apart from HEK293FT which were purchased from ThermoFisher.

Hypoxic incubations were carried out at 1% O_2_ and 5% CO_2_ in a humidified InvivO_2_ 400 hypoxic workstation (Baker-Ruskinn).

For lentiviral transduction HEK293T cells were co-transfected with lentiviral vector, pHR’-CMV-8.2ΔR and pMDG.2 plasmid prior to clearing viral supernatant and incubating with cells prior to selection with appropriate antibiotics or cell sorting via flow cytometry.

For transient transfections, DNA was mixed with serum-free medium (Opti-MEM, Gibco) and mixed with viafect (Promega) at a 3:1 ratio. Transfection mix was then added to cells for 24-48 hours before analysis.

### Plasmids

pLNT-HRE-JDG was a gift from Tristan R McKay. Tet-pLKO-puro was a gift from Dmitri Wiederschain (Addgene plasmid # 21915), shRNA inserts were annealed together and ligated into vector with AgeI and EcoRI restriction sites. Oligoes were shScr (top):

*CCGGCAACAAGATGAAGAGCACCAACTCGAGTTGGTGCTCTTCATCTTGTTGTT*

*TTTG. shSCR* (bottom): *AATTCAAAAACAACAAGATGAAGAGCACCAACTCGAGTTGGTGCTCTTCATCTT GTTG*. sh*COASY_1* (top): *CCGGCCTACCCAACACGCTGGTATTCTCGAGAATACCAGCGTGTTGGGTAGGT*

*TTTTG.* sh*COASY_1* (bottom): *AATTCAAAAACCTACCCAACACGCTGGTATTCTCGAGAATACCAGCGTGTTGGG TAGG.* sh*COASY_2* (top): *CCGGGCCACGTTTGAGGTTCTTGATCTCGAGATCAAGAACCTCAAACGTGGCTT*

*TTTG.* sh*COASY_2* (bottom):

*AATTCAAAAAGCCACGTTTGAGGTTCTTGATCTCGAGATCAAGAACCTCAAACG*

*TGGC* sh*COASY_3* (top):

*CCGGGCTGAAGATACTCACGGACATCTCGAGATGTCCGTGAGTATCTTCAGCTT*

*TTTG.* sh*COASY_3* (bottom):

*AATTCAAAAAGCTGAAGATACTCACGGACATCTCGAGATGTCCGTGAGTATCTT*

*CAGC.* PANK1β and *COASY* CDS were purchased from GenScript and subcloned into pCDH-EF1-FHC, which was a gift from Richard Wood (Addgene plasmid # 64874). HA-HIF1alpha-pcDNA3 was a gift from William Kaelin (Addgene plasmid # 18949). myc-BioID2-MCS was a gift from Kyle Roux (Addgene plasmid # 74223). HIF1A CDS was amplified from HA-HIF1alpha-pcDNA3 flanked with NotI restriction sites before being subcloned into myc-BioID2-MCS. Cassette expressing HIF1A-BioID2 was amplified by PCR flanked with Attb sites before being shuttled into pDONR221 and further cloned into pLX304-IRES-GFP (gift from William Kaelin) with gateway cloning. pcDNA3-UBFD1-Flag plasmid was purchased from GenScript and deletion mutants were generated using site-directed mutagenesis. For recombinant protein expression, the coding sequence corresponding to UBFD1-PH domain was amplified with primers forward: *TACTTCCAATCCAATGCCCCTCTCTGCAGGCAG* and reverse: *TTATCCACTTCCAATGTTAAAAATACTGCCATTTCCCCAG* and cloned into pCMSG7 using ligation independent cloning (LIC). PH domain cystine to alanine mutant was generated with site-directed mutagenesis.

### HRE-reporter screen

A549-HRE-Luciferase cells were transfected with siGENOME protein Kinase library (Horizon Biosciences), with INTERFERin (polyplus) transfection reagent at 50 nM. This library includes 720 SMARTpool siRNA targeting known kinases, pseudokinases and proteins with predicted kinase activity. Edge wells were filled with siRNAs targeting *HIF1A, HIF2A, PHD2* and scrambled control. 48 hours post transfection cells were maintained in hypoxia or normoxia for further 18 hours. Resazurin solution was added to a final concentration of 0.4 mg.mL^-1^ and cell viability was measured. Cells were lysed in passive lysis buffer (Promega) before measuring luciferase activity using Luciferase Assay substrate as per manufacturer’s instructions (Promega).

Luminescence values from luciferase activity were normalised to resazurin fluorescence readings. Z-scores were calculated for each kinase. siGENOME SMARTpool siRNAs against kinases with Z-scores ± 2 were purchased screened in validation screen and results were calculated as log2 fold change normalised luciferase activity over scrambled control.

### Immunoblot

Cells were lysed in RIPA buffer (50mM Tris-HCl (pH 8.0), 150 mM NaCl, 0.5% (w/v) deoxycholic acid, 0.1% (w/v) SDS & 1% (v/v) IGEPAL) before clearing lysate and determining protein concentration with Pierce BCA protein assay kit (ThermoFisher). Lysates were boiled in 5X SDS-PAGE sample buffer and electrophoresed on acrylamide gels, transferred onto PVDF membranes and immunoblotted with primary antibodies (See below). Anti-IgG horseradish peroxidase (Dako) and chemiluminescent detection (Immobilon, Millipore) were used to detect signal on Amersham Imager 600 (GE healthcare).

Antibodies used: HIF-1α (BD Biosciences, #610959), HIF-2α (Novus Biologicals, #NB100-122), HIF-1beta/ARNT D28F3 (Cell Signalling Technology, #5537S), CoAsy (abcam, ab129012), PHD2 (abcam, ab4561), β-actin (Sigma-Aldrich, #A1978), HIF-1α P564-OH (Cell Signalling Technology, #3434), Pank1 (Proteintech, #11768-1-AP), HA-Tag C29F4 (Cell Signalling Technology, #3724S), UBFD1 (Bethyl Laboratories, A305-143A), Flag/DYKDDDDK-Tag 9A3 (Cell Signalling Technology, #8146S), CoA (Gift from Ivan Gout), Flag M2 (Sigma-Aldrich, F1804), PSMD2 (Proteintech, 14748-1-AP), PSMC1 (Proteintech, 11196-1-AP), PSMA7 (Proteintech, 15219-1-AP).

### Immunoprecipitation and pulldowns

For co-immunoprecipitation of ectopically expressed proteins, cells were lysed with ice-cold NP-40 buffer (50 mM Tris (pH 8.0), 150 mM NaCl, 0.7% NP-40, 5% glycerol), lysate was cleared by centrifugation and incubated with pre-conjugated anti-HA beads (Pierce, #88837) for 1h at room temperature. Beads were washed with PBS-T (0.05% Tween) before eluting protein complexes with 1X SDS loading buffer diluted with NP-40 buffer.

To perform endogenous co-immunoprecipitations, protein G beads (Cytivia, #28953465) were pre-incubated with primary antibody (1 μg per IP) and 2% BSA before washing and adding cleared lysate for 1h at room temperature. Beads were washed with PBS-T (0.05% Tween) before eluting protein complexes with 1X SDS loading buffer diluted with NP-40 buffer.

Pulldowns with CoA-Agarose were performed by incubating CoA-Agarose beads (Sigma-Aldrich, C7013) or Sepharose control beads with NP-40 overnight at 4⁰C to hydrate and equilibrate the resin. Lysates or purified proteins were incubated for 1h at room temperature before washing with PBS-T (0.05% Tween) and eluting with 1X SDS loading buffer.

### Proximity labelling of HIF-1α interactome with BioID2

HEK293T cells expressing inducible *COASY* or *SCR* shRNA were transduced with pLX304-BioID2-HIF1A-IRES-GFP or pLX304-IRES-GFP and sorted to ensure equal GFP expression. RNAi knockdown was induced through treatment with doxycycline (25 ng.mL^-1^) for 48h before adding 50 μM Biotin for a further 18h. Cells were treated with MG132 (10 μM) for 6 hours before lysing cells in RIPA buffer. Biotinylated proteins were purified with streptavidin magnetic beads (Cytivia, #28985738). Beads were washed with: 2X RIPA, 1X KCl (1 M), 1X Na_2_CO_3_ (0.1M), 1X Urea (2 M in 10 mM Tris-HCl (pH8.0)) and 3X RIPA. Purified proteins were eluted by boiling at 95⁰C in FASP SDS buffer (2% SDS, 100 mM DTT, 50 mM Tris-HCl (pH7.5)).

### Filter Aided Sample Preparation (FASP) and LC-MS/MS analysis

FASP purification was performed as described previously^37^. Briefly, samples were diluted 10X with UA buffer (8 M urea, 100 mM Tris–HCl pH 8.8), and transferred to Vivacon Hydrosart 500 filters (30 kDa MW cutoff) before being concentrated by centrifugation. Proteins were alkylated by addition of 100 μL of 55 mM Iodoacetamide in UA buffer and incubation in the dark for 30 minutes, before being washed three times with 200 μL of UA buffer through cycles of concentration and reconstitution. This was followed by three washes with 100 μL of ABC buffer (40 mM ammonium bicarbonate). Proteins were then digested with 50 uL of trypsin (10 ng/μL) in ABC buffer overnight, in a 37⁰C humidified chamber. Peptides were then eluted by centrifugation. Two additional elutions were performed using 40 μL of ABC, followed by a final elution with 40 μl of 30% acetonitrile. The combined eluates were desalted using the Stage Tip procedure^38^ and recovered in A* buffer (0.1% TFA, 0.5% Acetic Acid, 2% Acetonitrile). LC–MS/MS analysis was performed on a Thermo Fisher Q Exactive-plus Orbitrap mass spectrometer coupled with a nanoflow ultimate 3000 RSL nano HPLC platform, as described previously^39^. Briefly, A* resuspended peptides were injected into the nanoflow HPLC and resolved at flow rate of 250 nl/min on an Easy-Spray 50 cm × 75 μm RSLC C18 column (Thermo Fisher), using a 123 min gradient of 3% to 35% of Buffer B (0.1% FA in Acetonitrile) against Buffer A (0.1% FA in LC–MS gradient water). The resolved peptides were infused into the MS by electrospray ionization (ESI), using a spray voltage of 1.95 kV, and a capillary temperature of 255°C. The MS was operated in data-dependent positive mode, with an MS1 scan at 70,000 resolution, followed by 15 MS2 scans at 17,500 resolution (top 15 methods). A 30-s dynamic exclusion was used to minimise repeat sequencing of the same peptides.

### Recombinant protein expression and purification

Recombinant proteins were expressed as before. Briefly, plasmids containing His-UBFD1-PH or His-UBFD1-PH-C→A were transformed into BL21 (DE3) competent cells (New England Biolabs, C2527H). Once liquid cultures were established, protein expression was induced by addition of 0.5 mM IPTG overnight at room temperature. Pellets were lysed with RIPA buffer and His-tagged proteins were isolated with HIS-Select Nickel magnetic beads (Sigma-Aldrich, H9914) before eluting with imidazole (300 mM Imidazole, 20 mM Tris-HCl (pH 8.0), 150 mM NaCl, 2.5 mM CaCl_2_).

### Thermal Shift assay

For each 50 μL reaction, 10 ug of His-UBFD1-PH was incubated with CoA or calcium pantothenate for 30 minutes at room temperature in a 50 mM NaCl and 50 mM Bis-Tris (pH 6.5) solution containing 10X Sypro Orange (Sigma-Aldrich, S5692). Samples were heated from 25⁰C to 95⁰C at a ramp rate of 1⁰C.min^-1^ and data acquired every degree on a Quantstudio 5 qRT-PCR machine (Applied Bioscience). Thermal shift assay curves were generated using TSA-CRAFT web server^40^ (https://bioserv.cbs.cnrs.fr/TSA_CRAFT/).

### Protein CoAlation

20 ug of recombinant protein was incubated with 1 mM CoA for 10 minutes at room temperature in a 50 mM Tris-HCl (pH 7.4) 150 mM NaCl buffer, prior to incubating with 300 μM H_2_O_2_ for a further 30 minutes. Reactions were stopped with the addition of 10 mM N-ethylmaleimide (NEM) and mixing with 5X SDS-sample buffer with or without β-mercaptoethanol.

### Bioinformatic Analysis

TCGA (Firehose legacy), METABRIC and CPTAC data was accessed via cbioportal^41^. DEPMAP data was accessed via the online portal^17^. Kaplan-Meier curves were generated using KMPlotter^42^.

### Colony formation Assays

Cells were plated at 250-1,000 cells per well, depending on cell line and cultured for 7-14 days in normoxia or hypoxia. Culture medium was changed every 2-3 days. At endpoint, colonies were fixed with 100% methanol, before being stained with 0.01% Crystal Violet (aq). Colonies were imaged (Amersham Imager 600) and counted using ImageJ.

### Orthotopic Xenograft Study

All animal procedures described in this project were approved by the UK Home office and designed with the principles of the NC3Rs and in adherence with ARRIVE guidelines. Seven-week-old NOD/SCID Gamma mice were purchased from Charles River and housed with food and water ad libitum, 6 animals were kept per cage. MDA-MB231 cells expressing indicated construct and H2B-mCherry were implanted into the mammary fat pad of mice (1 x 10^6^ cells in 50 μL of 1:1 PBS/Matrigel). Tumours were measured with callipers 3 times per week and the volume was calculated using the formula *V = (length*^2^ *x width)/2*. Two mice from v/o-FH group had tumour cell leakage during implantation and were therefore excluded from analysis. At endpoint, mice were culled, tumours and lungs were harvested and fixed in 10% neutral buffered formalin for downstream analysis.

### Immunohistochemistry

Formalin-fixed paraffin-embedded (FFPE) mouse tumours were sliced into 4 µm thick sections. Slides were baked for 2 hours at 60 °C prior to dewaxing. Afterwards, slides were put into xylene, 100% ethanol, and in 3% H2O2/methanol solution for 10 min for endogenous peroxidase inactivation. Antigen retrieval was performed using a 10 mM citrate buffer (pH 6) for 10 min at high power (700 W) in a microwave. Slides were blocked with goat serum and then incubated with Cleaved Caspase 3 primary antibody (1:100 overnight at 4⁰C, Cell Signalling Technology, #9661). The following day samples were incubated with ImmPress HRP Anti-rabbit polymer (Vector Laboratories, MP-7401) as per manufacturers instructions and developed with VIP chromogen prior to counterstaining with haematoxylin (Vector Laboratories, H-3502), dehydrating and mounting with DPX mounting medium. Slides were imaged with Nanozoomer S210 scanner (Hamamatsu). DPX mounting medium was cleared by soaking in xylene overnight, before rehydrating, performing antigen retrieval and blocking previous secondary antibody with AffiniPure Fab fragment Goat Anti0Rabbit IgG (H+L) (Jackson Immunology, # 111-007-003) prior to staining with Ki67 (1:100, 1 hour at room temperature, abcam, ab16667) and processing as described previously with cleaved caspase 3.

### Digital Pathology

All image analysis was performed using QuPath 0.6.0-rc3^43^. Immunohistochemical staining for Ki67 and Cleaved Caspase 3 (CC3) was assessed using a standardised pipeline.

For both markers, tissue detection was performed using the Pixel Classifier on whole sections of primary breast tumours. Nuclear segmentation was carried out using the InstaSeg Brightfield model^44^. H-score quantification was performed by defining DAB mean intensity classification thresholds.

**Ki67 Analysis:** Ki67-stained slides were deconvoluted using a custom H-DAB estimated colour deconvolution matrix:

setColorDeconvolutionStains(’{“Name”: “H-DAB estimated”, “Stain 1”:

”Hematoxylin”, “Values 1”: “0.72812 0.61682 0.29895”, “Stain 2”: “DAB”, “Values

2”: “0.46447 0.75875 0.4567”, “Background”: “166 156 186”}’)

Following nuclear segmentation, DAB mean intensity thresholds were set at 0.35, 0.6, and 0.8 to compute H-score quantification.

**CC3 Analysis:** CC3-stained slides were deconvoluted using the H-DAB default matrix:

setColorDeconvolutionStains(’{”Name”: “H-DAB default”, “Stain 1”:

”Hematoxylin”, “Values 1”: “0.65111 0.70119 0.29049”, “Stain 2”: “DAB”, “Values

2”: “0.26917 0.56824 0.77759”, “Background”: “255 255 255”}’)

Following segmentation, DAB mean intensity thresholds were set at 0.3, 0.35, and 0.4 for H-score quantification.

### Statistics

Data were normalised to relevant controls as required. All data is shown as mean ± SD from at least 3 biological replicates unless otherwise stated. Statistical analysis was conducted using GraphPad Prism 9.0, using the appropriate statistical test for the number of groups and type of data generated from the experiment. Unless otherwise stated, all tests performed are two-sided. Shapiro-Wilk test was performed to test for normality before determining parametric or non-parametric tests. For multiple comparisons post hoc tests were Kruskal-Wallis (One-way ANOVA) or Dunnett’s (Two-way ANOVA). Statistical significance is shown using the following nomenclature: ns p > 0.05, *p ≤ 0.05, **p ≤ 0.01, ***p ≤ 0.001, ****p ≤ 0.0001.

### Generative AI Usage Statement

In preparing this manuscript, generative AI tools—including QuillBot-Premium, ChatGPT 4.5(OpenAI), and Grammarly (1.109.2.0)—assisted with grammar, spelling, clarity, and text flow. These tools were used solely for linguistic refinement to improve readability and coherence, ensuring that the scientific content is communicated effectively. At no stage were AI tools used for idea generation, data analysis, result interpretation, scientific conclusions, or drafting of original scientific content.

All intellectual contributions, hypotheses, methodologies, experimental designs, data interpretation, and conclusions presented in this work are entirely the product of the author’s expertise and research. The use of AI was strictly limited to text editing functions and did not influence the integrity, novelty, or originality of the scientific findings.

This declaration is made in the interest of full transparency and adherence to ethical publishing standards, in accordance with Nature Press editorial policies on responsible AI usage.

## Data availability

The Mass spectrometry data generated during this study have been deposited to ProteomeXchange Consortium via the PRIDE partner repository with the dataset identifier (insert upon publication). Figures, siRNA screen results and list of proteins identified by BioID2-MS are available on Figshare (10.6084/m9.figshare.28566236). All other data related to this study are available upon request

## Acknowledgements

This research was supported by the Animal Technician Service “CRUK Animal Technician Service grant at Barts Cancer Institute (Core Award CTRQQR-2021\100004)”. This research was supported by the CRUK FLOW CYTOMETRY CORE SERVICE GRANT at Barts Cancer Institute (CTRQQR-2021\100004). This research was supported by the CRUK MICROSCOPY CORE SERVICE GRANT at Barts Cancer Institute (Core Award CTRQQR-2021\100004). The authors acknowledge the technical staff within the BCI Pathology core Facility at the Queen Mary University of London for technical support & assistance in this work. The authors would like to thank Barrie Peck and Andrew Finch for their helpful discussions with this work.

## Contributions

Paul Grevitt and Tyson V. Sharp designed and performed experiments. Oscar Maiques Carlos analysed tumour and lung sections. Martin Dodel and Faraz Mardakheh performed MS analysis and data processing. Michael Plevin performed structural analysis of UBFD1. HEK293T-Pank1β cells and anti-CoA antibody were provided by Ivan Gout who also provided technical advice throughout the project. Emanuela Gadaleta conducted analysis of an in-house patient cohort under the supervision of Claude Chelala and sample access was provided by Louise Jones. Kunal Shah, Katiuscia Bianchi and Valle Morales contributed with technical advice throughout this project. Tyson V. Sharp supervised and managed all research.

---

**Figure S1:**
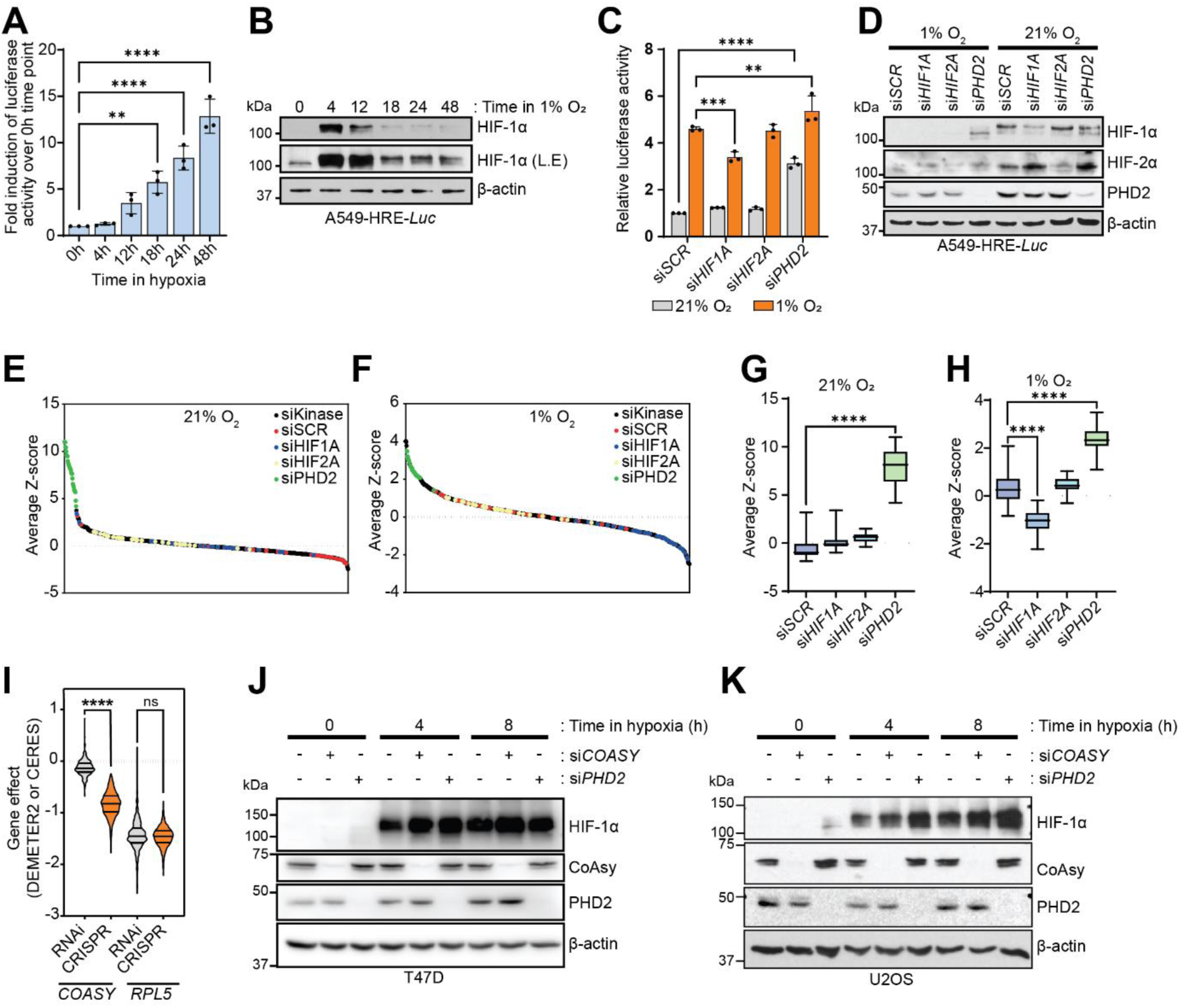
RNAi screen identifies Coenzyme A Synthase as a novel negative regulator of HIF1 signalling. **A)** Luciferase activity of A549-HRE-*Luc* cells cultured in hypoxia for indicated time (n=3). **B)** Western blot of A549-HRE-*Luc* cells for indicated time (n = 3). **C)** Luciferase activity of A549-HRE-*Luc* cells cultured in normoxia or hypoxia (18 h) after siRNA knockdown of *HIF1A*, *HIF2A* or *PHD2* (n = 3). **D)** Western blot of A549-HRE-*Luc* cells cultured in normoxia or hypoxia (18 h) after siRNA knockdown of *HIF1A*, *HIF2A* or *PHD2* (n = 3). **E-F)** Average Z-score from primary screen in normoxia (**E**) and hypoxia (**F**) (n = 2). **G-H)** Positive and negative controls from primary screen (n = 2). **I)** Gene essentiality scores from CRISPR (Ceres) or RNAi screens (DEMETER2) from DEPMAP database. **J)** Western blot of T47D cells with *COASY* knocked down via siRNA in normoxia or hypoxia (n = 3). **K)** Western blot of U2OS cells with *COASY* knocked down via siRNA in normoxia or hypoxia (n = 3). * < 0.05, ** p < 0.01, *** p < 0.001, **** p < 0.0001

**Figure S2:**
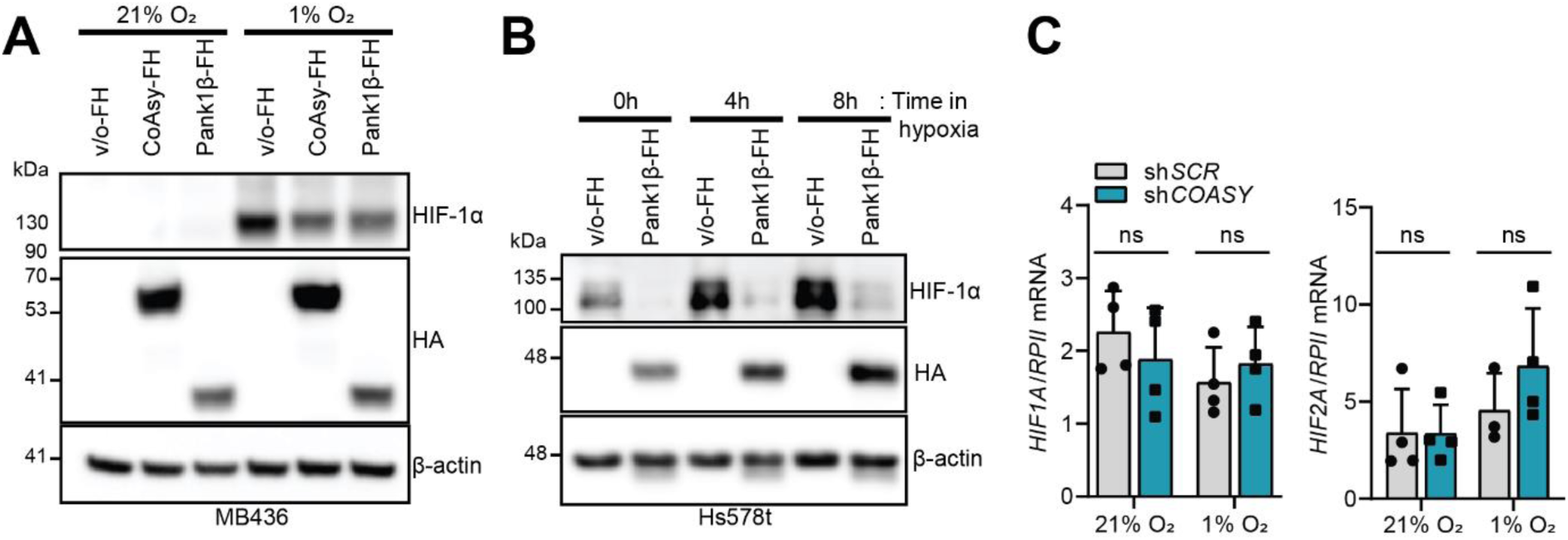
Coenzyme A biosynthesis is coupled with HIF signalling. **A)** Western blot of MB436 cells overexpressing CoAsy-FlagHA or Pank1β-FlagHA in normoxia or hypoxia (4h). **B)** Hs578t cells overexpressing Pank1β-FlagHA in normoxia or hypoxia. **C)** *HIF1A* and *HIF2A* mRNA expression in A549 sh*SCR* or sh*COASY* cells in normoxia and hypoxia (n = 4).

**Figure S3:**
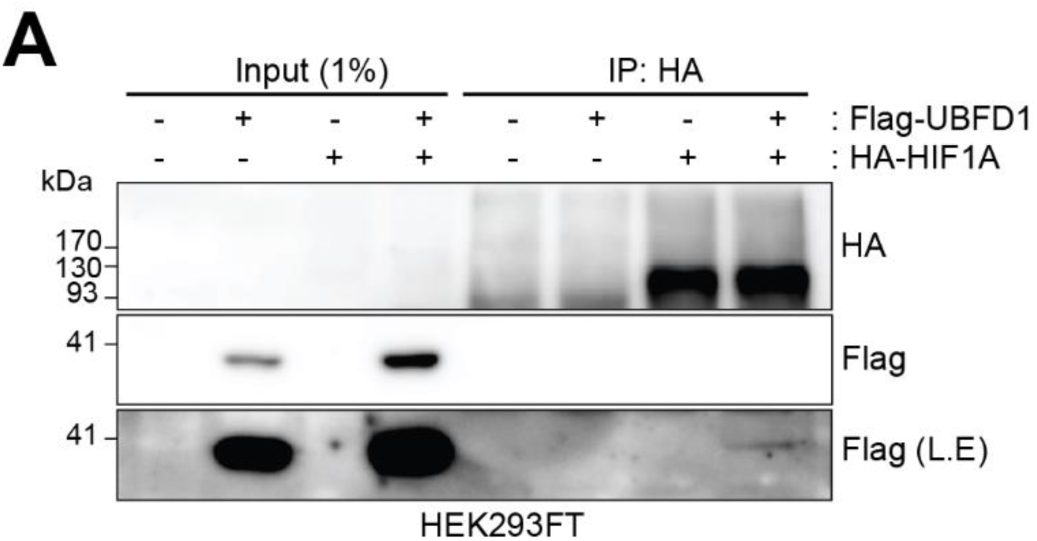
Proximity labelling-MS identifies UBFD1 as a CoAsy dependent interactor of HIF-1α. **A**) Overexpressed co-immunoprecipitation of HA-HIF1A with Flag-UBFD1 (n = 5). L.E = long exposure.

**Figure S4:**
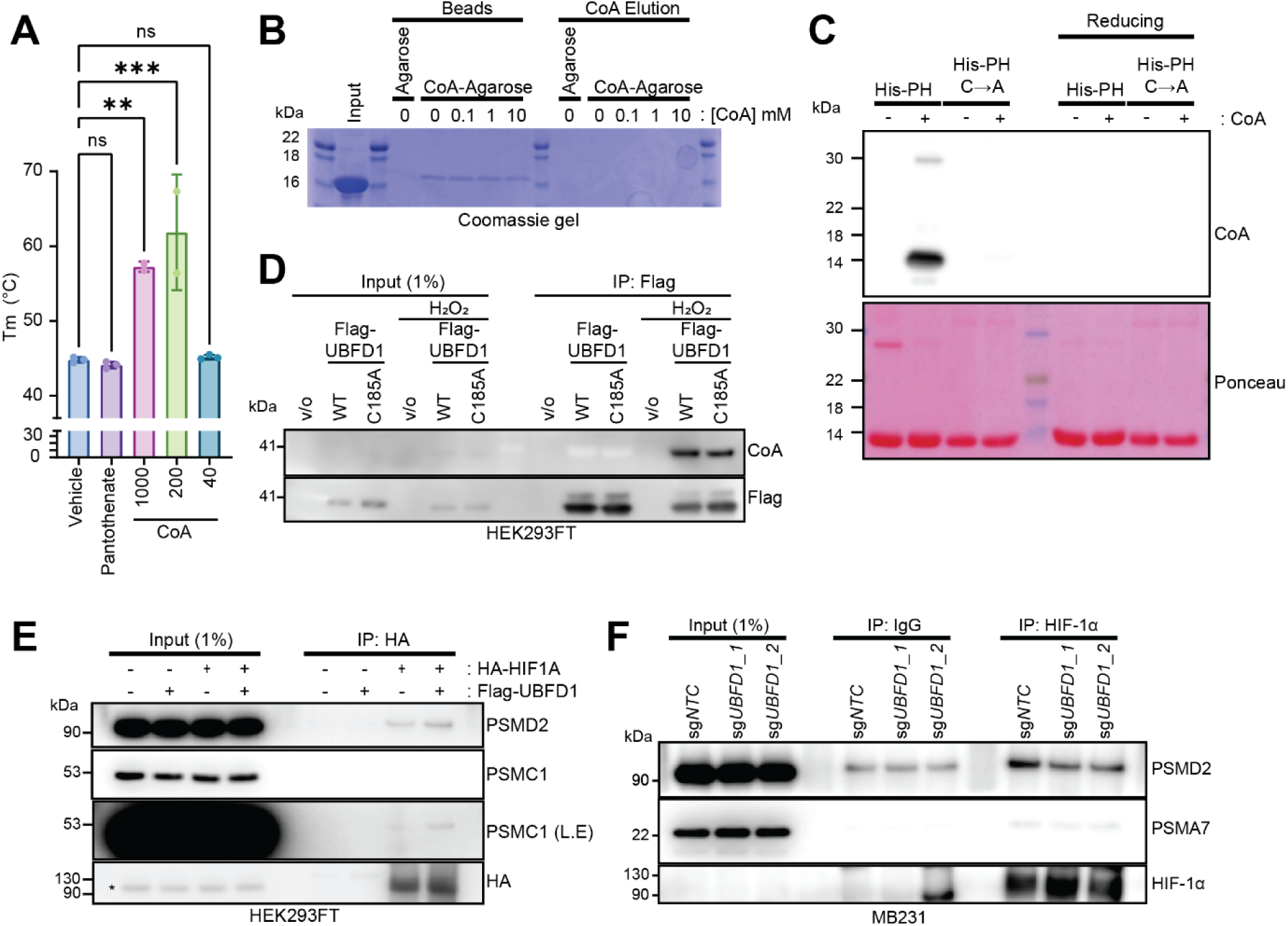
UBFD1 is CoAlated which controls association with HIF-1α. **A)** Melting temperature of His-UBFD1-PH domain incubated with indicated concentration of CoA as determined by thermal shift assay (n = 1). **B)** Pulldown of His-UBFD1-PH domain with CoA agarose beads, and eluate with increasing amounts of CoA (n = 2). **C)** Western blot of in vitro CoAlation assay of His-UBFD1-PH domain and cysteine to alanine mutant (n = 3). **D)** Western blot of HEK293FT cells overexpressing Flag-UBFD1 or C185A mutant treated with H2O2 (n=2). **E)** Co-immunoprecipitation of HA-HIF1A and proteasome components in cells overexpressing Flag-UBFD1 (n = 2) *indicates non-specific band. **F)** Co-immunoprecipitation of HIF-1α and proteasome components in MB231 cells with UBFD1 knocked out via CRISPR-Cas9 (n = 3). L.E = long exposure. * < 0.05, ** p < 0.01, *** p < 0.001.

**Figure S5:**
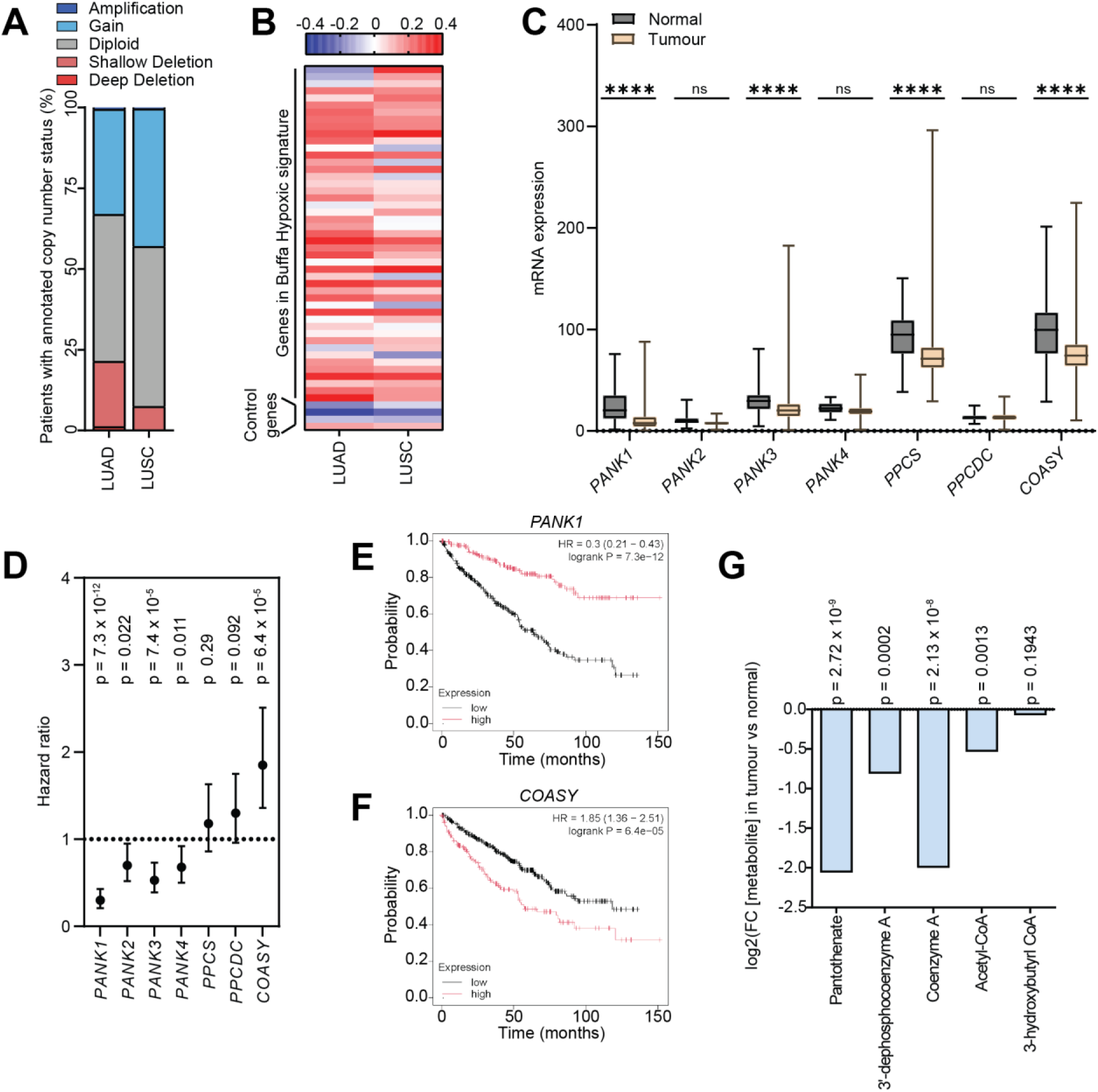
COASY as a biomarker in lung cancer and clear cell renal cell carcinoma (ccRCC). **A)** Putative copy number alterations of *COASY* in LUAD and LUSC from the TCGA dataset. **B)** Heatmap of spearman correlation coefficients between *COASY* and genes from Buffa hypoxic signature in LUAD and LUSC. **C)** mRNA expression of genes in CoA biosynthesis pathway in clear cell renal cell carcinoma. **D)** Hazard ratios of ccRCC patients’ overall survival stratified by genes in CoA biosynthesis. **E-F)** Kaplan-Meier plots for patients stratified by *PANK1* (**E**) or *COASY* (**F**). **G)** Metabolite abundance in tumour compared to normal tissue from Li et al. *Nature* 2014^29^. * < 0.05, ** p < 0.01, *** p < 0.001, **** p < 0.0001

**Figure S6:**
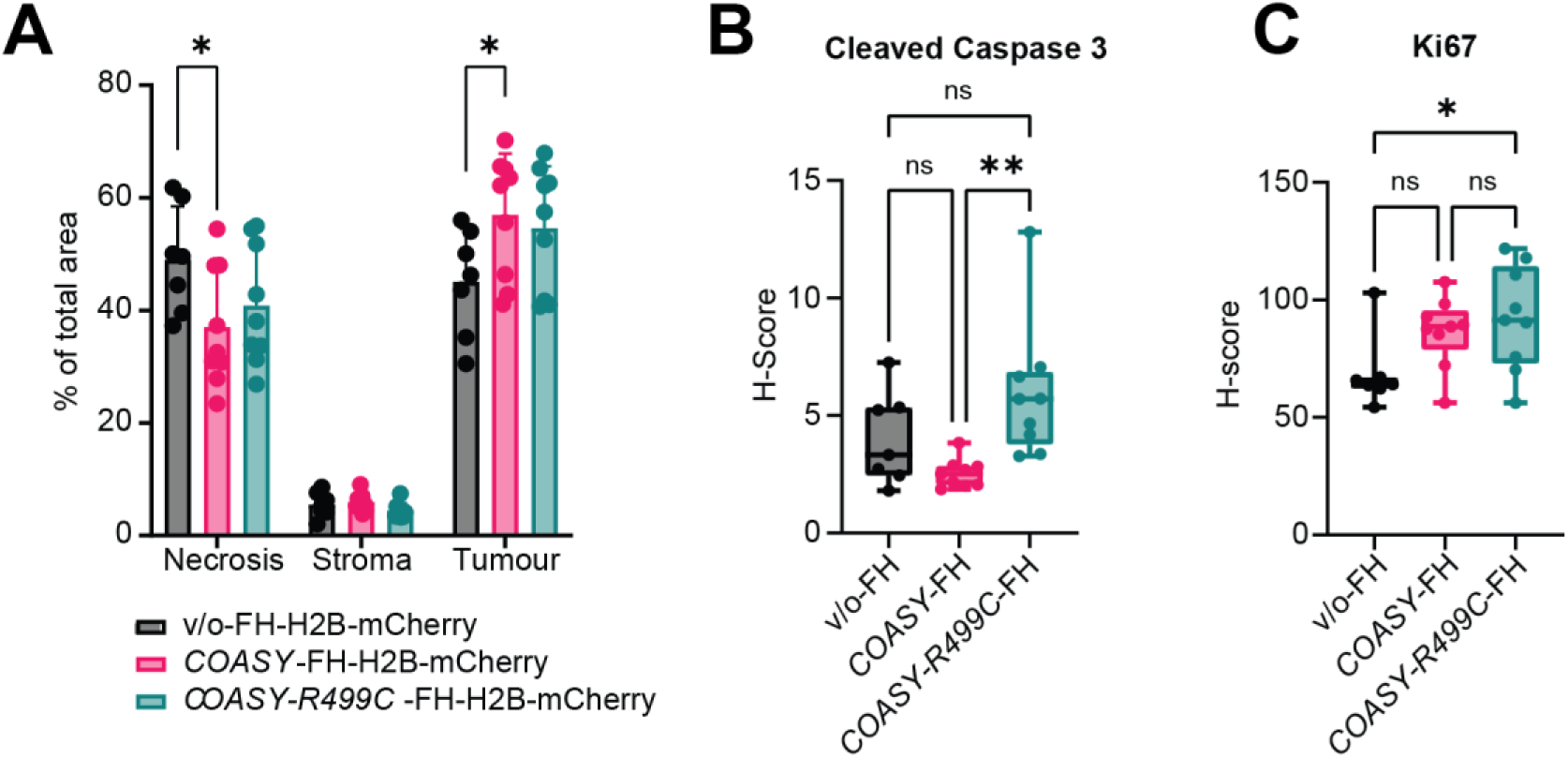
CoAsy overexpression does not alter primary tumour growth in TNBC models. **A)** Quantification of necrosis, stroma and tumour cells in MB231-FH orthotopic tumours from H&E stains. **B-C)** H-Scores of markers for apoptosis (Cleaved Caspase 3, **B**) and proliferation (Ki67, **C**) from orthotopic tumours. * < 0.05, ** p < 0.01.

